# Disentangling folding from energetic traps in simulations of disordered proteins

**DOI:** 10.1101/2024.11.06.622270

**Authors:** Jeffrey M. Lotthammer, Alex S. Holehouse

## Abstract

Protein conformational heterogeneity plays an essential role in a myriad of different biological processes. Extensive conformational heterogeneity is especially characteristic of intrinsically disordered proteins and protein regions (collectively IDRs), which lack a well-defined three-dimensional structure and instead rapidly exchange between a diverse ensemble of configurations. An emerging paradigm recognizes that the conformational biases encoded in IDR ensembles can play a central role in their biological function, necessitating understanding these sequence-ensemble relations. All-atom simulations have provided critical insight into our modern understanding of the solution behavior of IDRs. However, effectively exploring the accessible conformational space associated with large, heterogeneous ensembles is challenging. In particular, identifying poorly sampled or energetically trapped regions of disordered proteins in simulations often relies on qualitative assessment based on visual inspection of simulations and/or analysis data. These approaches, while convenient, run the risk of masking poorly-sampled simulations. In this work, we present an algorithm for quantifying per-residue local conformational heterogeneity in protein simulations. Our work builds on prior work and compares the similarity between backbone dihedral angle distributions generated from molecular simulations in a limiting polymer model and across independent all-atom simulations. In this regime, the polymer model serves as a statistical reference model for extensive conformational heterogeneity in a real chain. Quantitative comparisons of probability vectors generated from these simulations reveal the extent of conformational sampling in a simulation, enabling us to distinguish between situations in which protein regions are well-sampled, poorly-sampled, or folded. To demonstrate the effectiveness of this approach, we apply our algorithm to several toy, synthetic, and biological systems. Accurately assessing local conformational sampling in simulations of IDRs will help better quantify new enhanced sampling methods, ensure force field comparisons are equivalent, and provide confidence that conclusions drawn from simulations are robust.

## INTRODUCTION

Over the last few decades, all-atom simulations of proteins have served as a “computational microscope,” enabling the interrogation of protein structure and dynamics at atomistic resolution^1–3^. All-atom simulations have been increasingly used to generate new hypotheses and interpret experimental data, enabling an iterative cycle where experiments can guide simulations and simulations guide experiments ^4^. Despite remarkable progress, there remain ongoing challenges in the application and interpretation of all-atom simulations, especially in the context of large biomolecular systems^5^.

Broadly defined, two main challenges exist for protein simulations. Firstly, despite continual advancements over the last several decades, there are still limitations in the accuracy and throughput of force fields^6–14^. Secondly, even on modern hardware, sampling the vast conformational landscapes of proteins is challenging^15–21^. Over the years, all-atom force fields have been consistently improving, and the development of these force fields remains an area of active research^6–8,10–14,22–24^. Moreover, a myriad of tools and techniques have been developed and leveraged to enhance conformational sampling^15,25–35^. However, despite remarkable progress in both areas, computationally sampling the energy landscape for proteins remains a challenge.

Energy landscape theory is a conceptual framework for understanding protein conformational space ^36–39^. In this model, the probability of occupying a configuration at any point within the high-dimensional “phase space” is proportional to the free energy of that configuration. For proteins that fold, the underlying landscape is often depicted as a high-dimensional funneled space, whereby the bottom of the hypothetical “folding funnel” corresponds to the native state (**Fig. 1A**) ^40–44^. While conformational fluctuations and protein dynamics in and around the native state can influence and even dictate molecular function, these fluctuations typically retain a (near) native-like conformation, reducing the potential collection of isoenergetic configurations dramatically from the universe of all possible configurations given the many degrees of freedom in a modestly-sized polypeptide chain ^45,46^. In contrast, for proteins that do not fold, no such native state exists, opening the set of isoenergetic (and hence equally likely) states to be vastly increased compared to their folded counterparts (**Fig. 1B**). Consequently, while exploration of configurational space is certainly not trivial for folded domains, this problem is dramatically exacerbated in the case of intrinsically disordered proteins and protein regions (collectively referred to as IDRs).

**Figure 1:**
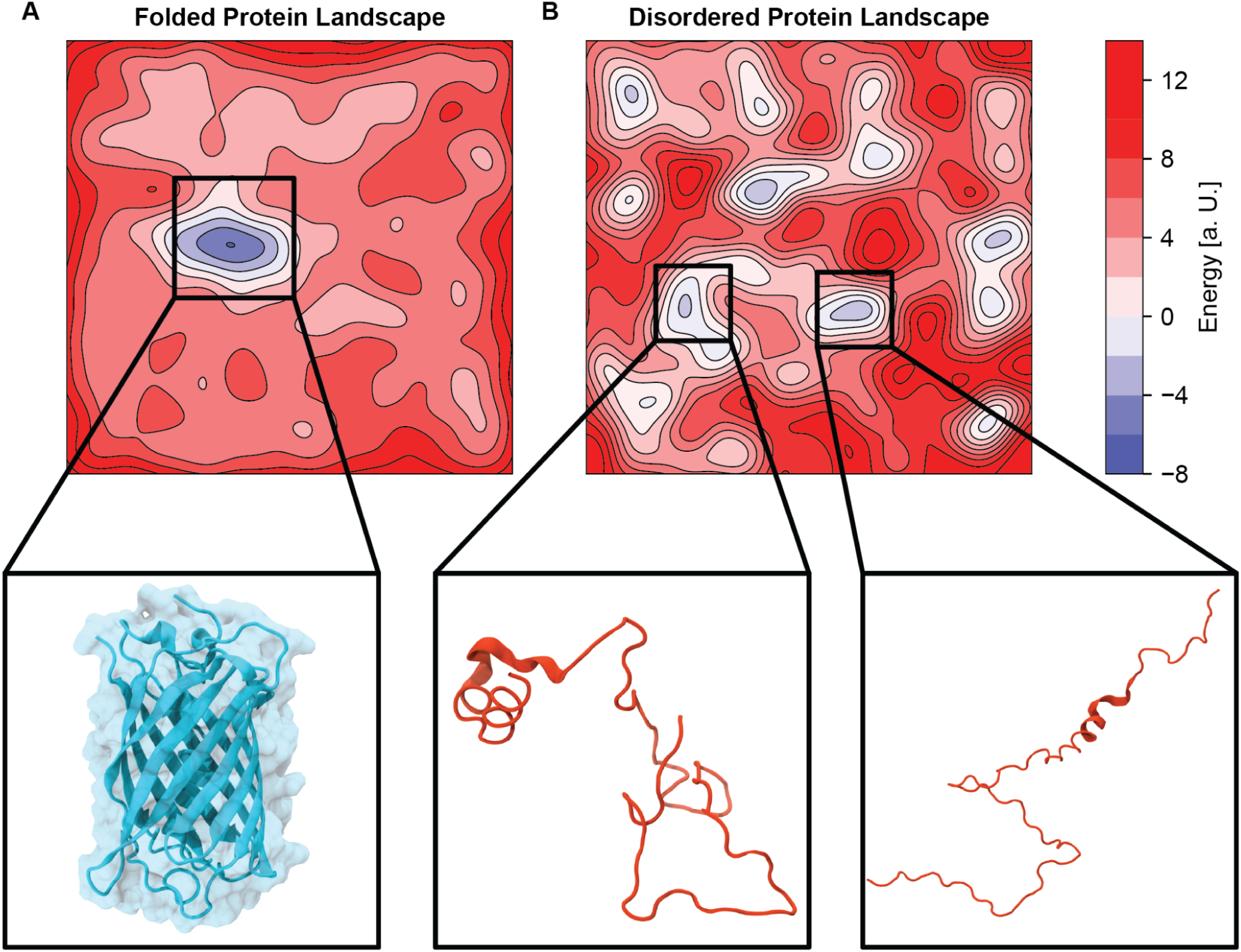
Hypothetical two-dimensional energetic landscape contour plots for A) a folded protein and B) a disordered protein. A gradient of blue to white to red is used to represent regions of lot to high energy. The folded protein has a singular clear minimum whereas the disordered protein possesses many distinct low energy basins. Axes and unit scales are arbitrary. The insets represent exemplar low energy configurations for each protein.

The challenge associated with conformational sampling for IDRs can be considered in terms of two related but distinct problems. First, a large configurational space necessitates exploration to identify energetically accessible conformations that are structurally distinct from one another. Second, simulations must be able to identify local minima and reasonably report on their contributions to the underlying landscape while also “escaping” these meta-stable configurations in the pursuit of sampling many distinct local minima. While these two challenges are well-known to the simulation community, it is often appealing to address the former using longer simulations. While this may be an effective strategy in certain situations, if escape from local minima is vanishingly unlikely, doubling or tripling simulation length will not impact the variety of distinct configurations seen. As such, a wide range of enhanced sampling approaches have been developed to improve the ability of simulations to explore configurational space^15,25–35^. Despite this, we generally lack systematic approaches to assess sampling quality in simulations, especially in the context of IDRs.

Assessing the adequacy of conformational sampling is a classic problem in the field of molecular simulations^16,17^. Visual inspection is a powerful and necessary check that can quickly identify very poor sampling, i.e., where no conformational rearrangement occurs at all. However, visually identifying "good" sampling is much more challenging.. Many heuristics and analyses have been developed in the context of folded domains, where a defined 3D reference structure offers a structural fiducial marker from which conformational fluctuations can be compared^5,16^. Given IDRs lack a stable folded state, approaches that rely on a defined reference structure are generally inapplicable for assessing sampling in IDRs. The importance of reasonable conformational sampling when drawing conclusions from simulations of IDRs cannot be overstated. In the worst-case scenario, simulations run for many microseconds may, in fact, only sample a small number of distinct conformations, making generalizable conclusions impossible. More problematically still, within a single IDR, distinct subregions may be sampled to different extents, hampering a single overall score from being useful or interpretable. Taken together, this gap in knowledge hinders our understanding of the conformational behavior of IDRs.

In this work, we build on prior work and apply limiting polymer models as reference distributions from which local conformational sampling can be compared. Inspired by the previous work of Lyle *et al.*^47^, we develop and implement an algorithmic approach to quantitatively assess local conformational sampling in IDRs. Our approach focuses on assessing the collective accessible distributions of dihedral angles in the polypeptide chain, and compares observed distributions to expected distributions from an all-atom resolution excluded volume-based simulation as a statistical reference model for well-sampled IDR conformational ensembles. Moreover, by comparing distribution obtained from multiple independent simulations, our approaches enable us to distinguish between the scenario in which IDR consistently adopts a specific and limited set of dihedrals (i.e., local or global folding) or adopts distinct but limited sets of dihedrals in different simulations (energetically-trapped). The resulting algorithmic pipeline PENGUIN (**<u>P</u>**ipeline for **<u>E</u>**valuating co**<u>N</u>**formational hetero**<u>G</u>**eneity in **<u>U</u>**nstructured prote**<u>IN</u>**s), is implemented in the SOURSOP simulation analysis package^48^ and can be used with all-atom molecular dynamics and Monte Carlo simulations.

## METHODS

### 1.1 Overview

Previous work has focused on developing physically rigorous tools to more efficiently sample conformational space via either thermodynamically perturbative enhanced sampling or statistically biased adaptive sampling algorithms such as umbrella sampling, replica exchange, Progress-Guided Index Sampling, FAST, parallel tempering, and weighted ensemble simulations (to name but a few) ^15,16,27,29–32,35,49,50^. In contrast, this work asks a different question: Namely, given a simulated ensemble, how widely are we exploring conformational space? Previous work by Lyle *et al*. devised a quantitative measure to assess conformational heterogeneity in disordered protein ensembles^47^. This approach focuses on assessing global ensemble properties, enabling changes in conformational heterogeneity to be assessed as a function of sequence or under different conditions, such as temperature or salt. More recent work by González-Delgado *et al*. developed an approach for the comparison of multiple IDR ensembles on a local and global level with specific applications for convergence analysis^51^. These approaches offer a formal route to quantify conformational heterogeneity with potential applications for understanding changes in configurational entropy upon binding, changes in solution environment, and in the context of refinement to experimental data.

In contrast, PENGUIN was developed for the more practical application of rapidly and easily assessing if conclusions from a simulation of a disordered protein are reliable. The underlying methodology applies a multi-ensemble comparison approach, necessitating computational complexities compatible with many independent comparisons. We choose to focus on local regions of the ensemble with residue level granularity and ask the question: how well sampled and/or how reproducible are our simulations via comparisons to statistical reference models? Importantly, we leverage an inter- and intra-trajectory analysis approach, which enables us to distinguish between finding a minimum on the energy landscape and staying there (i.e., “folding”) versus becoming trapped in one of many possible local minima on the landscape. A key feature of PENGUIN is our use of the atomistic Excluded Volume (EV) ensemble to define a well-sampled reference distribution of a real chain with unambiguous mapping to the true ensemble (**Fig. 2A**).

**Figure 2:**
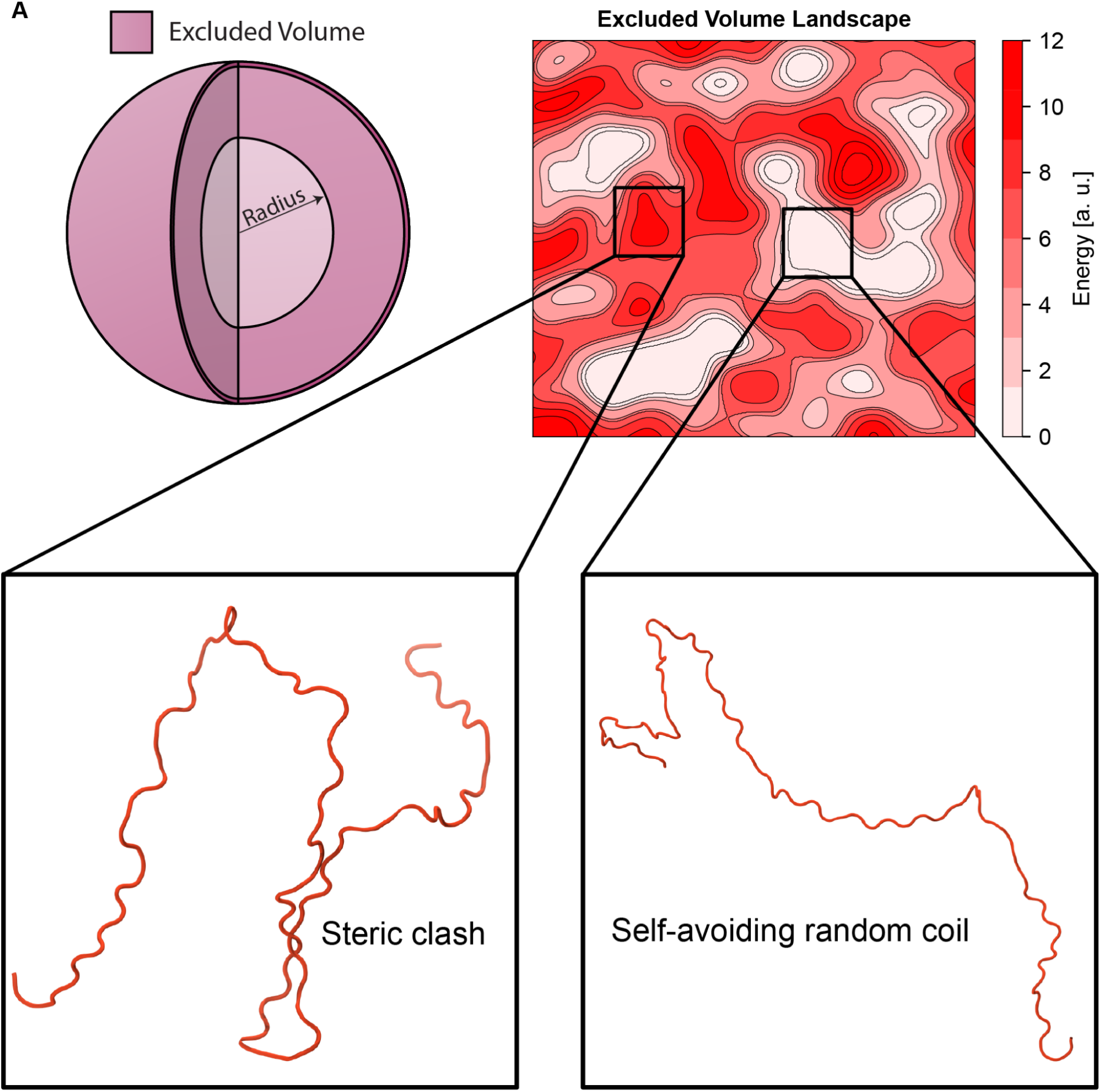
The excluded volume limit is a reference model for biopolymers. **A)** Three-dimensional cross section of the excluded volume model limit between two spheres. The excluded volume is determined by the radius of the circle. **B)** Two-dimensional energetic landscape contour plot in the excluded volume limit for the same hypothetical disordered protein in Figure 1B. A gradient of blue to white to red is used to represent regions of low to high energy. Notably, the landscape of the excluded volume limit has lower energetic barriers than the hypothetical landscape in Figure 1B. The inset depicts a representative conformation of the biopolymer chain in the excluded volume limit. The chain is free to orient itself in any direction; however, the residues are not allowed to overlap with other residues in the chain.

The EV limit reflects a specific class of simulation in which all attractive and repulsive terms in the underlying force field are ignored except for bond lengths/angles and the repulsive component of the short-range (e.g., Lennard-Jones) potential (**Fig. 2B; left inset**) ^52–54^. This is in contrast to what we refer to as the Full Hamiltonian (FH) limit, whereby all terms in the force field are used. In the EV limit, the polymer chain is non-overlapping, yet no contributions of electrostatics, solvent, or local non-bonded effects bias the resulting ensemble (**Fig 2B, right inset**). Ensembles generated from EV simulations follow statistics for self-avoid random coils, yet unlike analytical models, also offer specific local and global structural distributions in the limit of the sampling obtained from the simulation. This feature makes EV simulations useful for direct comparison with FH simulations in that identical simulation analysis procedures can be applied to EV ensembles, as can ensembles generated using all the force field components. This is in contrast to analytical models, whereby comparisons are limited to observables where relevant parameters can be derived^55–57^. Consequently, EV simulations have been used extensively as a polymer reference state to which structural properties for disordered protein ensembles can be compared ^54,55,58–62^.

Given that polymer models are rooted in the statistical distributions of flexible chains, they are particularly well suited to describe the conformational behavior of unfolded and disordered proteins ^52,62–65^. However, biological molecules are not ideal polymers. Instead, the underlying physicochemical properties of the amino acid sequence create deviations from ideal behavior; biopolymers are neither random coils nor excluded volume models^62,66^. Indeed, biopolymers have preferential attractive and repulsive interactions that modulate conformational heterogeneity (**Fig 1B, 2B**)^62,67–72^. Nevertheless, the excluded volume model is a powerful reference as it serves as a sequence-specific theoretical upper bound on the configurational entropy for a real chain. This coarsening of the underlying energy landscape also alleviates some of the sampling difficulties found in FH simulations of disordered proteins, as many barriers are reduced (**Fig 1B, 2B**). To assess conformational sampling, numerical simulations in both the FH and EV limits are performed, and residue level conformational heterogeneity is computed for each of the simulated trajectories. By scoring the similarity in conformational heterogeneity between these two models, we provide a quantitative metric for assessing if a given residue or region of sequence is robustly sampled within the limits of the available number of conformers. This unbiased scheme allows for the assessment of local conformational heterogeneity in simulations of disordered proteins.

The remainder of this paper is organized as follows: we describe the algorithmic workflow associated with PENGUIN and perform a series of analyses to illustrate its utility. We first demonstrate the feasibility of this approach by testing it on a representative toy system where we expect efficient sampling to be tractable. Then, as a control, we encode poor sampling in our simulations and demonstrate the metric is able to recapitulate the expected behavior for a poorly sampled sequence region. Further, we examine synthetic constructs where we would expect energetically trapped states due to nonspecific hydrophobic collapse and demonstrate that our metric shows that this sequence is poorly sampled. Finally, we examine several previously published IDR simulation datasets.

### 1.2 The PENGUIN Algorithm

Our algorithmic pipeline, PENGUIN, has two primary goals. First, we wish to quantify how well local subregions are sampling conformational space for a single simulation. Second, if we observe that a region appears ‘poorly sampled’, we must be able to delineate between the scenario in which independent simulations yield different energetically trapped configurations (i.e., many local traps) compared to a situation in which independent simulations repeatedly arrive at the same low-energy state (i.e., protein folding). To enable this, the PENGUIN pipeline can be summarized as follows:

(1) For the disordered protein of interest, perform one Full Hamiltonian atomistic simulation starting from a random initial configuration.
(2) Perform a complementary simulation in the EV limit or take advantage of precomputed distributions from excluded volume tripeptide simulations (see methods).
(3) Compute a dihedral angle probability mass function for each residue from each simulation in the EV and FH limit.
(4) For each residue, compute the Hellinger distance – also referred to as Jeffreys distance – (*Equation 1*) to score the similarities between the dihedral angle distributions for each residue between the respective ensembles. The 1D Hellinger distance for two discrete probability distributions given by *P* = (*p*_1_, … , *p_k_* ) and *Q* = (*q*_1_, … , *q_k_*) can be computed as follows:

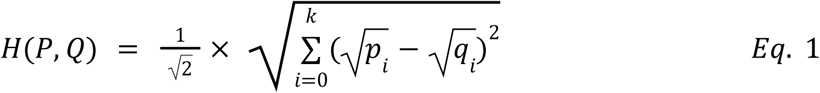

where P is a dihedral angle (e.g., φ or ψ for a specific residue) probability vector for the FH ensemble, Q is the dihedral angle probability vector for the EV ensemble for the corresponding residue, and angle i is an index that ranges from 1 to k, where k is the total number of discretized bins, and p_i_ or q_i_ is the probability of finding the angle in question in bin i for the FH ensemble (p_i_) or EV ensemble (q_i_).

The 2D Hellinger distance (e.g., as would be used for simultaneous comparison of a 2D distribution in φ and ψ dihedral space) is given by:

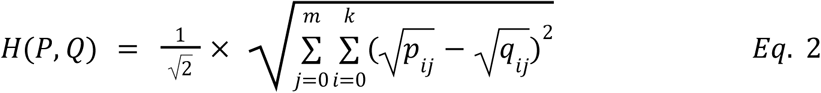

Where the terms have the same meaning, except there are two distinct dimensions (j and i) over which the distance is calculated.

(5) Repeat steps 1-4 for a total of N pairs of replicate simulations.
(6) Assess locally conformational sampling by comparing the Hellinger distance between EV and FH simulations at each residue across many independent replicas. If this distance is large for any given residue, it implies that FH simulations are locally trapped.
(7) To delineate the differences between many distinct energetically trapped configurations vs. the same energetically trapped configuration being identified in multiple independent simulations (e.g., protein folding), perform a complementary analysis using the Hellinger distance to compare all-to-all trajectory values on the FH simulations.

The core components of this algorithm are summarized graphically in **Figure 3**.

**Figure 3:**
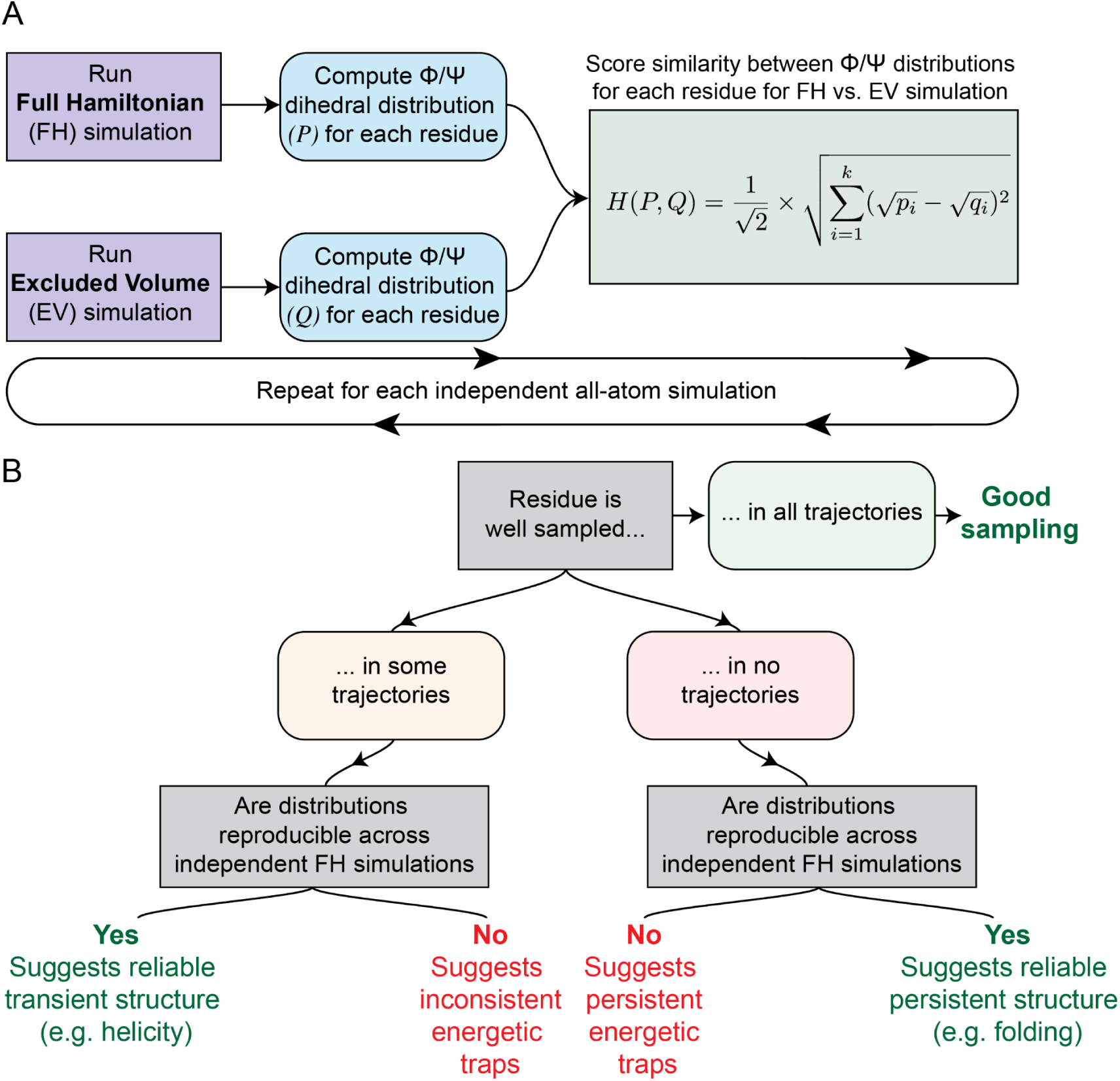
An algorithmic pipeline for assessing and quantifying disordered protein conformational heterogeneity. **A)** A flow chart depicting the pipeline for quality assessment. First, perform two sets of simulations with N replicates in both a physics-based potential and in the excluded volume limit. Then, compute the dihedral angle probability distributions in each limit. Score their similarity and repeat for each set of replicates. **B)** Comparison amongst a set of replicate simulations is used to ascertain whether a particular residue is widely sampling conformational space, folding or partially folding, or energetically trapped undergoing nonspecific hydrophobic collapse.

The Hellinger distance metric is used to compare the similarities of two probability distributions because it is symmetric, nonnegative, and satisfies both the identity of indiscernibles, and the triangle inequality. This metric is bounded between 0 and 1 (i.e., 0 ≤ *H*(*P*, *Q*) ≤ 1). If the distributions are similar, then the Hellinger distance will be near 0 (exactly 0 for identical distributions). In contrast, if the distributions are dissimilar, the Hellinger distance will be closer to 1. While various metrics for measuring the distance between two distributions exist, we use the Hellinger distance for its ease of implementation, ability to easily handle instances of disjoint supports, and - importantly - fast computation, enabling rapid comparisons of IDR ensembles at scale.

PENGUIN is implemented in the SOURSOP simulation analysis package ^48^ within the **`sssampling`** module (https://github.com/holehouse-lab/soursop). The associated functions are documented under the associated **sssampling** documentation on the SOURSOP ReadTheDocs page (https://soursop.readthedocs.io/en/latest/modules/sssampling.html).

### 1.3 Monte Carlo Simulations

All Monte Carlo simulations first reported in this paper were performed using the ABSINTH-OPLS/AA implicit solvent model with the CAMPARI simulation engine^22,73–75^. The monovalent ion parameters derived by Mao et al. were used^76^. Unless otherwise stated, all standard CAMPARI degrees of freedom (backbone and sidechain dihedral angles, ion position, relative protein position) were sampled. For the FS-peptide with fixed residues, a CAMPARI freeze file was used to avoid sampling backbone or sidechain dihedrals for residues 3-10. The CAMPARI keyfiles, sequence files, and freeze files used for each Monte Carlo simulation can be found in the associated directory at the Holehouse lab Supporting Data GitHub repository under https://github.com/holehouse-lab/supportingdata/. Simulation data for Ash1 monte carlo simulations were obtained from ^61^.

A summary of the simulations performed as part of this study is included in **Table 1**:

**Table 1.** Table of simulations details and associated figures reported in this study.

| Name and associated figure | #reps | Equilibration (steps) | Production (steps) | Output frequency | Conformers per simulation | Total conformers |
| --- | --- | --- | --- | --- | --- | --- |
| FS-peptide: Fig 4. | 20 | 5 000 000 | 125 000 000 | 40 000 | 3000 | 60 000 |
| FS-peptide (fixed): Fig 5. | 20 | 5 000 000 | 125 000 000 | 40 000 | 3000 | 60 000 |
| (GV)<sub>20</sub>-(PGSK)<sub>10</sub>: Fig 6 | 20 | 20 000 000 | 200 000 000 | 50 000 | 3600 | 72 000 |
| Ash1: Fig 7. | 15 | 2 000 000 | 32 000 000 | 5 000 | 6000 | 90 000 |
| GCN4 (150K - 400K): -<br>Fig S3 | 15 | 5 000 000 | 35 000 000 | 10 000 | 3000 | 45 000 |
| Tripeptides | 1 | 0 | 250 000 | 250 | 1000 | 1000 |

### 1.4 Precomputed EV distributions

To make complex and resource-intensive EV simulations more accessible, we developed an interface that approximates EV simulations using precomputed dihedral angle distributions from tripeptide simulations. These simulations, which included NME and ACE capping groups, were performed with the same settings as our other Monte Carlo simulations. We precomputed dihedral angle distributions for all 20 standard amino acids in the context of each of the 20 amino acids, resulting in 8000 different tripeptide simulations. This comprehensive set of context-dependent dihedral angle distributions forms the basis of our approximation method.

We use predefined mappings to retrieve the appropriate dihedral angle distributions based on the residue and its context. For a given protein sequence, our algorithm shreds the sequence into a list of 3-mers, followed by a 2-mer or 1-mer if the sequence length is not divisible by 3. For the 1-mers or 2-mers, we pad the sequence with alanines and extract the corresponding tripeptide phi or psi angles as needed. To generate dihedral angles that match the length of the simulations, we sample from these distributions using a histogram-based approach, which involves creating a histogram of the precomputed angles, calculating the cumulative distribution function (CDF), and using inverse transform sampling to generate new angles that follow the same distribution for the appropriate length, e.g., # of frames, for the simulation. The sampled angles are then replicated for the specified number of trajectories, providing a set of context-aware, approximated dihedral angles for each residue in the sequence.

This approach allows us to efficiently approximate the excluded volume dihedral angle distributions for any protein sequence, based on the precomputed tripeptide simulations performed in the excluded volume limit. By doing so, we provide a computationally efficient route to obtain sequence specific dihedral angle distributions in the absence of full-length simulations performed in the polymer limiting model.

### 1.5 1D vs. 2D Hellinger distance analysis

Computation of the Hellinger distances requires the comparison of two probability distributions. For implementation simplicity, we compute the dihedral angle probability distributions by discretizing the dihedral angle space into discrete bins and comparing the probabilities of occupying these bins for both the inter and intra ensemble comparisons. We chose to use a bin width of 15 degrees to discretize the dihedral angle space. This decision was made to match the choice of move set we use in our campari simulations for dihedral angle perturbations, although this is a free parameter to modulate in our implementation. We note that while there is some sensitivity to the bin width parameter at extreme values, the analysis is relatively insensitive to the discretization provided the bin width does not become too granular or envelop many different rotameric states. If the bin width is too granular, subtle differences in distributions will be exacerbated in the distance calculation due to finite data. Therefore, we recommend a bin width of 15 degrees which we set as default.

## RESULTS

### 3.1 Visualizing sampling quality with PENGUIN

A key feature of any analysis approach is that it should make it easy to draw general conclusions from the resulting data. After investigating several possible routes, we settled on a consistent four-panel format for visualizing PENGUIN-derived assessments of sampling quality. In all four panels, the x-axis reports on the per amino acid position, while the y-axis reports distinct information relating to the sampling quality.

The top left panel reports on the per-residue average (black) and per-trajectory (red) Hellinger distances between Excluded Volume (EV) and Full Hamiltonian (FH) simulations for a specific dihedral distribution (or 2D dihedral distribution) (e.g. **Fig. 4B**). If FH and EV dihedral distributions are highly similar, values here will approach zero. If EV and FH dihedral distributions are very different, values here will approach 1. If values here approach 0, this is a necessary and sufficient condition for good sampling, while if values approach 1, this is a necessary but not sufficient condition for bad sampling, but does indicate specific dihedrals are “far” from a fully-flexible chain. The spread across independent simulations here is indicative of sampling quality. If independent FH simulations cluster tightly around the average, this indicates that independent simulations are equivalently far from the EV ensemble, a necessary but not sufficient condition for good sampling. On the other hand, if independent FH simulations show a variety of different distances to EV simulations, this implies substantial simulation-to-simulation heterogeneity, which is necessary and sufficient for at the very least insufficient and potentially poor sampling.

**Figure 4:**
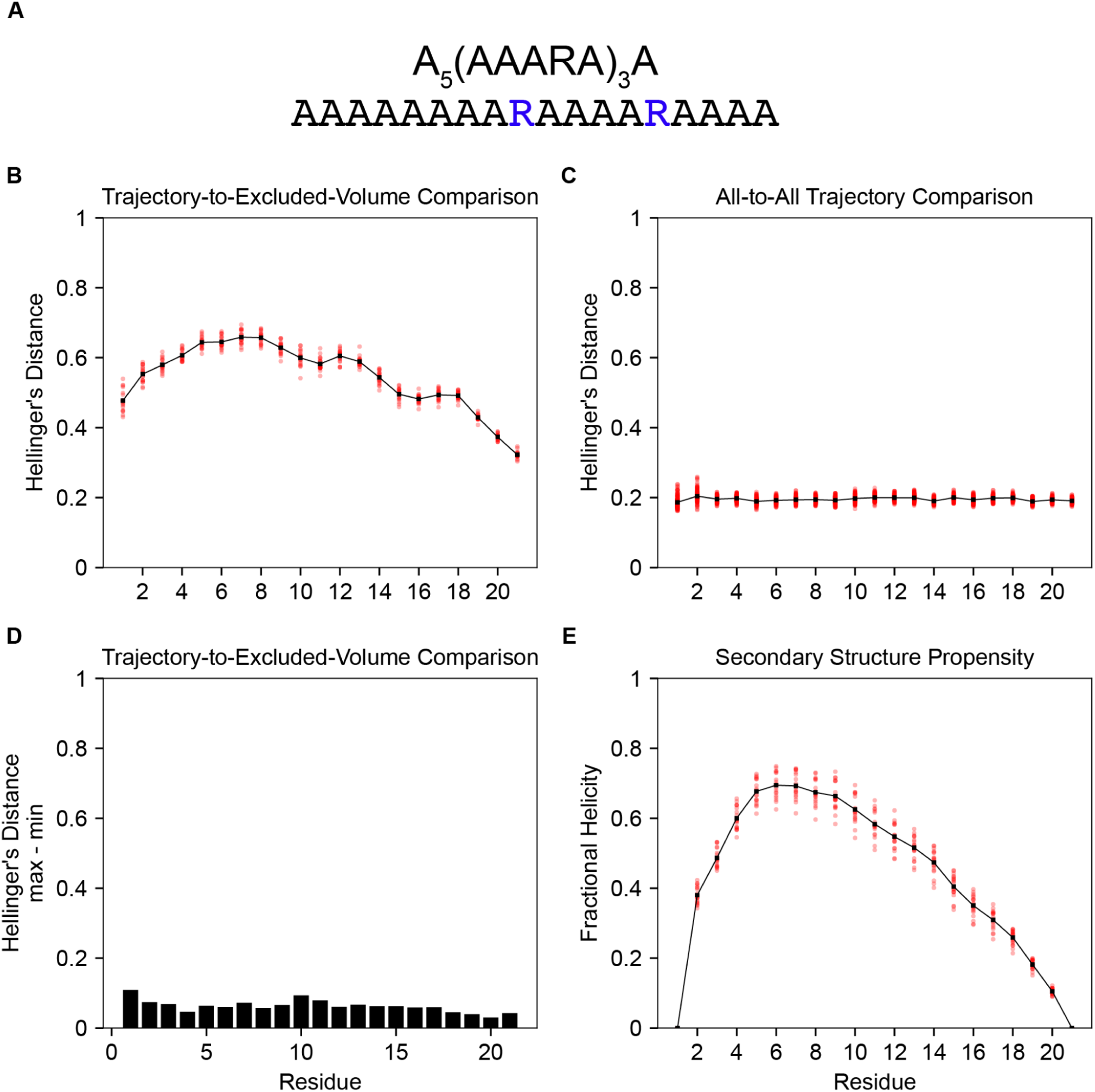
FS-peptide as a model system for assessing local conformational heterogeneity. **A)** The simulated sequence **B)** Per-residue Hellinger distances between the phi dihedral angle distribution of the simulated ensemble and the excluded volume ensemble. **C)** An all-to-all simulated ensemble comparison of the per-residue phi dihedral Hellinger distance. **D)** The maximum minus the minimum Hellinger distance for each residue from the distribution in B. **E)** The per-residue fractional helicity. In each panel, the transparent red points represent the raw data whereas the black line represents the mean between replicates.

The top right panel reports on the average (black) and per-trajectory (red) Hellinger distances between all independent FH simulations (e.g. **Fig. 4C**). In the limit of good sampling, independent simulations should be similar to one another, with a Hellinger distance approaching 0 and all red points clustering tightly around the average. Such a result is necessary and sufficient to conclude that the sampling is good. However, if sampling is poor, independent simulations may be trapped in distinct conformational states, yielding large variations in inter-simulation distance with independent simulations showing a wide variety, such that large variations and a higher average distance are necessary and sufficient to conclude poor sampling across many independent trajectories. This enables one to distinguish between a situation where sampling is very unlike the EV simulations and independent replicas are highly similar, indicating transient or persistent local or global folding.

The bottom left is an alternative representation of the information in the top left and reports on the spread of the Hellinger distances between the EV and FH simulations (e.g. **Fig. 4D**). If this is small, it is consistent with and indicative of good sampling. If this is large, it suggests *at least* one of the trajectories is likely poorly sampled. This can be useful for highlighting rare, poorly sampled trajectories that are lost on average but may substantially bias conclusions from ensemble analysis.

The bottom right reports on the average (black) and per-trajectory (red) local helicity for each residue, as assessed by the DSSP algorithm (e.g. **Fig. 4E**). This is a complementary (albeit distinct) type of analysis that can help explain and correlate results seen in the other three panels.

Over the following analyses, we apply figures with these four panels as a consistent approach to make comparisons between systems that are well or poorly sampled.

### 3.2 FS-peptide as an exemplar well-sampled simulation

We first turned to FS-peptide to test the PENGUIN’s ability to detect locally well-sampled regions in conformational space (**Fig. 4A**). The FS-peptide has been used for decades as a model system for exploring protein folding and stability^13,74,77,78^. We quantitatively assessed the difference between dihedral angle distributions by comparing the per-residue conformational heterogeneity of FS-peptide in the FH simulations and EV simulations. While our FH simulations deviate from the EV reference model, the dihedral angle distributions are highly reproducible across twenty independent replicas starting from distinct configurations (**Fig. 4B**). In addition to the insensitivity to starting configuration, the reproducibility of the dihedral angle distributions suggests that the observed deviations from the excluded volume limit, as measured by intermediate Hellinger distance values, are a consequence of the underlying physicochemical properties of the amino acid sequence. Moreover, an inter-trajectory comparison between the simulations performed for the FH simulations reveals that the residue level dihedral angle distributions are reproducibly observed (**Fig 4C**). As the Hellinger distance can be used to identify local sequence regions where there may be poor sampling, the statistical spread of the distribution of Hellinger distances is useful for quickly inspecting sampling quality across *n* replicate simulations, where *n* is the number of simulations performed. Therefore, to identify local sequence regions where at least one trajectory may exhibit poor sampling, we compute the per-residue statistical range of the dihedral angle Hellinger distances (**Fig. 4D**). For a well-sampled residue or sequence region, we would expect the statistical range of the Hellinger distance to be small. Our analysis reveals that all Hellinger distances observed for the well-sampled FS-peptide system are near zero. Consistent with the observations that these simulations deviate from the excluded volume limit, our simulations agree with prior studies and suggest the formation of transient helicity in FS-peptide (**Fig. 4E**).

### 3.3 Locally-frozen regions as facsimiles of folded domains

Next, we investigated an artificially “folded” system in which an eight-residue stretch (residues 3-10) of the FS-peptide is frozen (**Fig. 5A**). This setup means all simulations have an identical but entirely unsampled subregion, mimicking an extreme scenario in which a specific subregion reproducibly folds into a specific conformation. The resulting analysis highlights how PENGUIN enables the easy identification of poorly-sampled but foldable regions.

**Figure 5:**
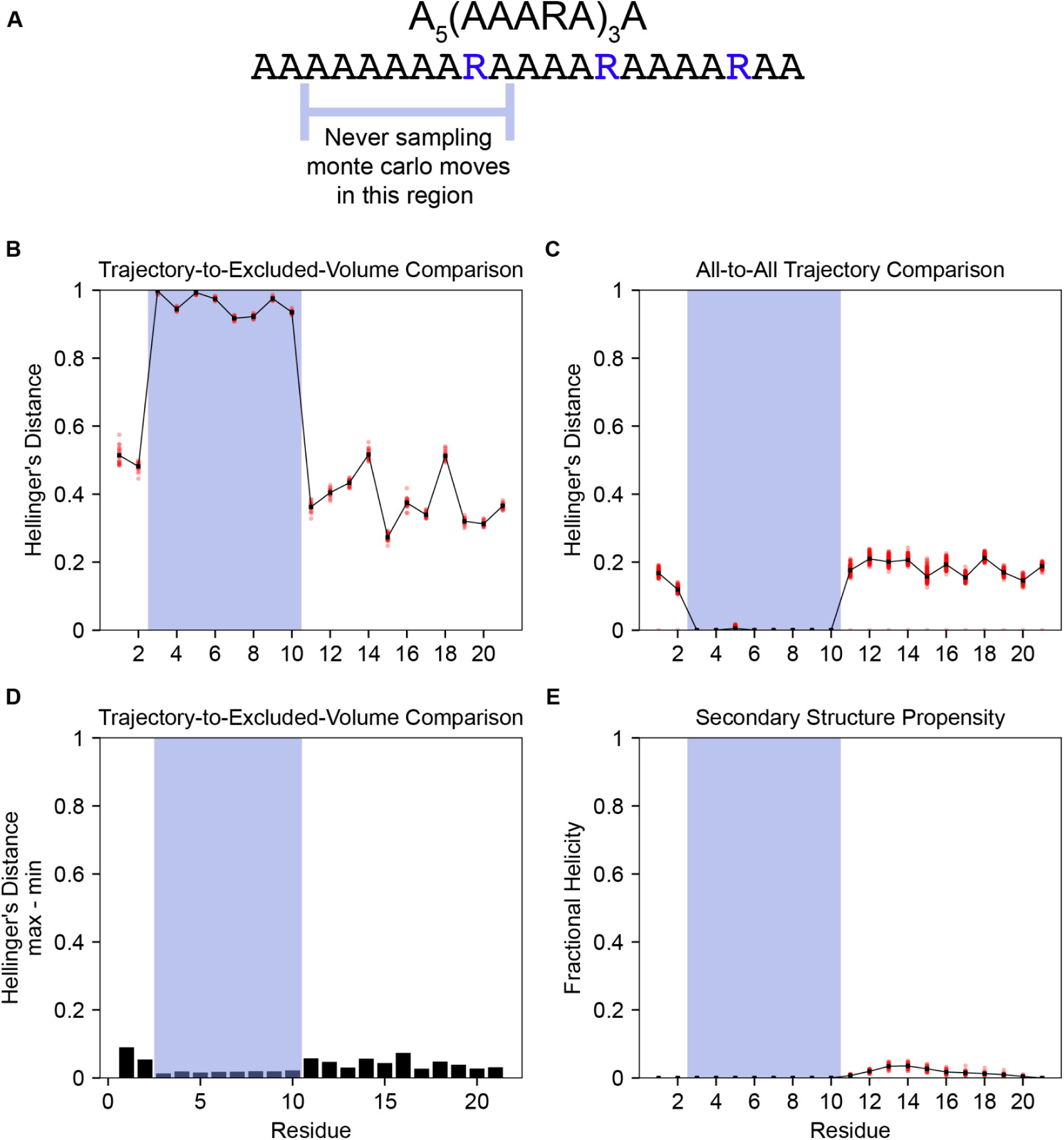
A fixed region of FS-peptide encodes poor local conformational sampling. **A)** The simulated sequence. The blue line beneath the sequence indicates the range of residues (3 to 10) that are fixed in this set of simulations.**B)** Per-residue Hellinger distances between the phi dihedral angle distribution of the simulated ensemble and the excluded volume ensemble. **C)** An all-to-all simulated ensemble comparison of the per-residue phi dihedral Hellinger distance. **D)** The maximum minus the minimum Hellinger distance for each residue from the distribution in **A. E)** The per-residue fractional helicity. In each panel, the transparent red points represent the raw data whereas the black line represents the mean between replicates. The amino acid sequence of FS-peptide is shown below. The corresponding fixed region is highlighted in blue in each panel.

Analysis of these systems confirmed that the frozen subsequence appears poorly sampled, as indicated by the Hellingers distances near 1 (**Fig. 5B**). However, importantly, the all-vs-all comparison between independent FH simulations revealed a Hilinger distance of ∼0 for the frozen residues, indicating very little inter-trajectory variation in these regions, consistent with a locally folded region (**Fig. 5C**). Finally, the per-residue max-min of the Hellinger distance between FH and EV simulations suggests that, despite large differences in the extent of sampling, the variation in replica data is extremely low across both frozen and flexible regions (**Fig. 5D**). Notably, however, the formation of fractional helicity for this set of simulations is markedly lower than in the simulation that is freely sampling conformational space (**Fig 4D, 5E**).

### 3.4 Poor sampling is clearly identifiable with PENGUIN

We next sought to determine whether we can differentiate heterogeneous and poorly sampled regions (i.e. regions trapped in energetic minima) from homogeneous and poorly-sampled regions (i.e. regions that fold). To this end, we designed an 80-residue low-complexity diblock copolymeric disordered sequence (**Fig. S1**) with an N-terminal hydrophobic region made of (GV)_20_ and a C-terminal charged and proline region made of (PGSK)_10_ (**Fig. 6A**). We reasoned, at least in our simulations, the hydrophobic N-terminus would undergo non-specific collapse becoming trapped in local energetic minima, while the C-terminal proline/charged rich block would remain highly expanded and well-sampled^61,71,79–81^.

**Figure 6:**
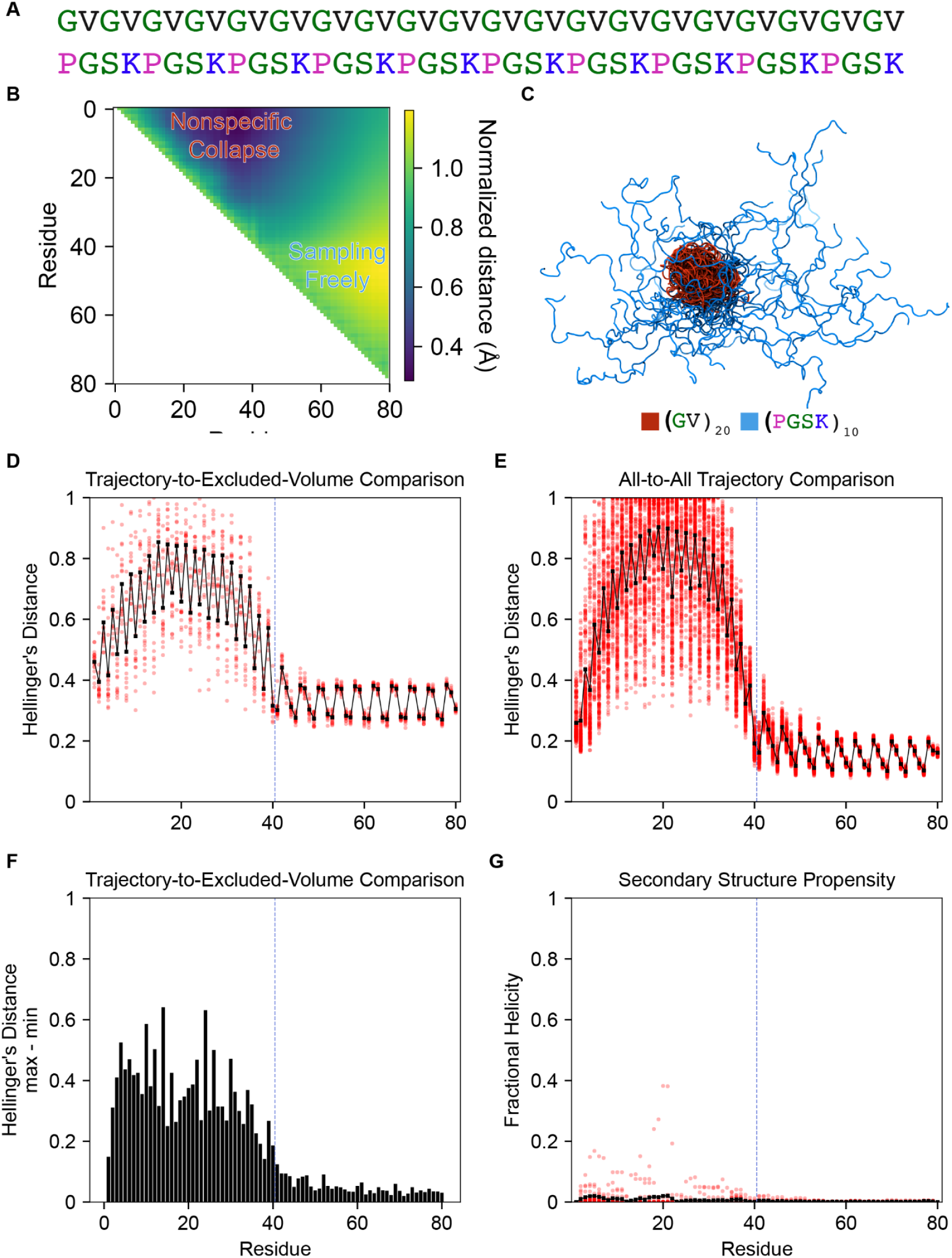
A hydrophobic patch in the synthetic sequence (GV)_20_-(PGSK)_10_ poorly samples conformational space. **A)** The simulated sequence **B)** An average inter-residue distance scaling map for the (GV)_20_-(PGSK)_10_ ensemble. Regions that are more collapsed than expected are shown in blue and regions that are more expanded are shown in yellow. Distance maps were normalized by comparison to EV simulations. Data shown is an average across n=10 replicate simulations. **C**) Molecular rendering of 100 representative conformations illustrating the (GV)_20_ region (red) is undergoing a nonspecific hydrophobic collapse whereas the (PGSK)_10_ region is sampling more freely. **D)** Per-residue Hellinger distances between the phi dihedral angle distribution of the simulated ensemble and the excluded volume ensemble. **E)** An all-to-all simulated ensemble comparison of the per-residue phi dihedral Hellinger distance. **F)** The maximum minus the minimum Hellinger distance for each residue from the distribution in A. **G)** The per-residue fractional helicity. In each panel, the transparent red points represent the raw data whereas the black line represents the mean between replicates. The blue dashed line delineates the region where the sequence transitions from GV to PGSK repeats.

Simulations of this synthetic sequence confirmed our expectations, with scaling maps of the sequence revealing two distinct subregions, a compact N-terminal region and an expanded C-terminal region (**Fig. 6B,C**). Moreover, the (GV)_20_ region (residues 1-40) appear poorly sampled, as suggested by the comparison to the EV limit in (**Fig 6D**). In addition to showing average values with a Hellinger distance far from EV simulations, independent trajectories showed a wide variety of distances from the EV simulations, indicating distinct dihedral distributions for different simulations. This inference is further confirmed by the all-to-all trajectory comparison, which revealed extensive heterogeneity in dihedral angle distributions across independent FH simulations (**Fig 6E**). Finally, the statistical range (max–min) of the Hellinger distances computed between independent FH replicas and EV simulation in the (GV)_20_ region (residues 1-40) further support the conclusion that the (GV)_20_ region in this synthetic sequence is poorly sampled.

In contrast, the proline-rich and positively-charged (PGSK)_10_ region (residues 40-80) shows robust sampling by all measures, indicating this region exists in similar, well-sampled conformational ensembles across each independent FH simulation (**Fig. 6**). This toy example exemplifies how sampling quality in a single polypeptide can be dramatically influenced by amino acid composition, and even within the same physical chain, some regions may be extremely well-sampled while others virtually not at all.

### 3.6 PENGUIN in the wild - examining ensembles of naturally-occurring sequences

Having examined several toy systems to calibrate our expectations and verify that PENGUIN enables us to disentangle between limits of good and poor sampling, we next sought to investigate how previously published ensembles fared when analyzed using PENGUIN. We focussed primarily on our own prior work, in part because if we retrospectively identified limitations in sampling, we felt comfortable that the original authors would not be influenced by this disclosure.

We first considered the now well-studied 83-residue disordered region from the yeast transcription factor Ash1 (residues 420-500; **Fig. 7A**)^61^. Mirroring simulation parameters used originally, PENGUIN suggests that by-and-large all-atom simulations of Ash1 are reasonably well-sampled (**Fig. 7B,C,D,E**). This dovetails with previous work demonstrating excellent agreement with Small Angle X-ray Scattering (SAXS) data for Ash1, as well as reasonable agreement with local secondary structure inferences. Of note, two small subregions (residues 40-50 and residues 50-60) appear somewhat less-well sampled than the remainder of the chain. It’s worth nothing that while residues 50-60 do show small amounts of residual helicity, residues 40-50 show none, and residues 10-20 which also show residual helicity are extremely well-sampled.

**Figure 7:**
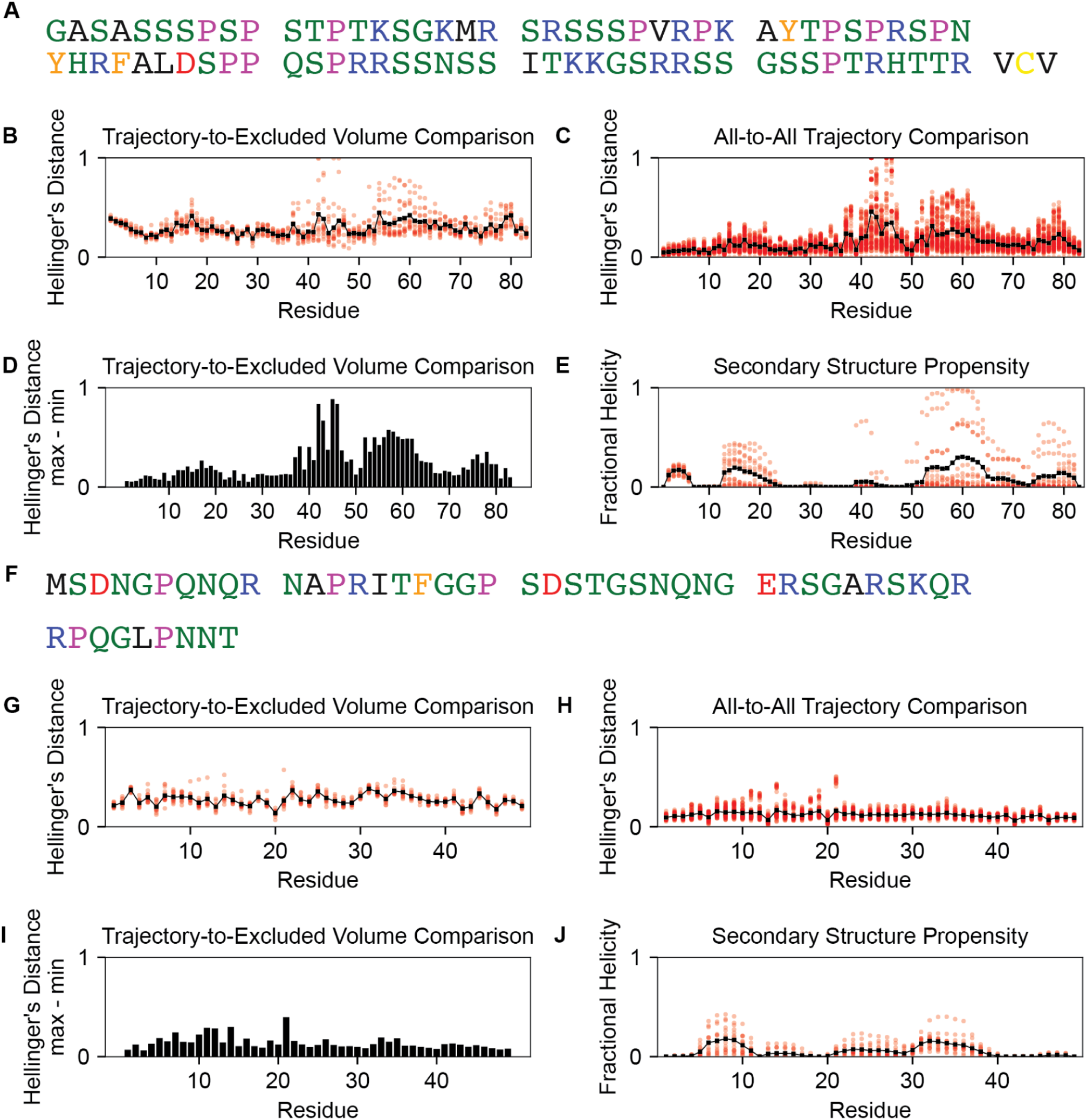
Extensive Monte Carlo simulations of biological systems reveal efficient exploration of conformational space. **A)** The sequence of Ash1^420–500^ with an N-terminal glycine-alanine dipeptide (herein referred to simply as Ash1). **B)** Per-residue Hellinger distances between the phi dihedral angle distribution of the simulated ensemble and the excluded volume ensemble for Ash1. **C)** An all-to-all simulated ensemble comparison of the per-residue dihedral Hellinger distance for Ash1. **D)** The maximum minus the minimum Hellinger distance for each residue from the distribution in **B. E)** The per-residue fractional helicity. In each panel, the transparent red points represent the raw data whereas the black line represents the mean between replicates. **F)** The sequence of the n-terminal domain (NTD) of the SARS-CoV-2 nucleocapsid protein **G)** Per-residue Hellinger distances between the phi dihedral angle distribution of the simulated ensemble and the excluded volume ensemble for NTD. **H)** An all-to-all simulated ensemble comparison of the per-residue phi dihedral Hellinger distance for the NTD of SARS CoV 2’s nucleocapsid protein. **I)** The maximum minus the minimum Hellinger distance for each residue from the distribution in **G. J)** The per-residue fractional helicity. In each panel, the transparent red points represent the raw data whereas the black line represents the mean between replicates.

We next considered the N-terminal IDR (NTD) from the SARS CoV-2 nucleocapsid (N) protein^82^ (**Fig. 7F**). We re-examined previously performed simulations of the NTD in isolation. Encouragingly, these simulations report extremely good local convergence across the entire chain, with near EV-like ensembles and very limited variation across independent trajectories (**Fig. 7G,H,IJ**). As such, we are able to conclude that – within the limits of forcefield accuracy – our conclusions from these simulations are unbiased by limited sampling.

### DISCUSSION & CONCLUSION

Assessing sampling quality and convergence in molecular simulations of proteins is crucial for ensuring the reliability and accuracy of the simulation results. Consequently, there is a rich history of both methodologies to improve conformational sampling as well as assess conformational sampling in these simulations. Ensuring high-quality sampling lends confidence that the simulation adequately explores protein conformational space and thus that any resulting analyses are representative, reflecting the true behavior of the protein under the modeled conditions. However, any simulation quality assessment approach will inevitably be blind to regions of conformational space that were never explored during the simulation. This limitation arises because the assessment is confined to the sampled data, meaning it cannot account for conformational states that were not observed. This problem particularly manifests itself in the analysis of disordered proteins due to their extensive structural heterogeneity and highly isoenergetic conformational landscapes.

A critical yet often underappreciated component of IDR ensembles is the sheer vastness of their conformational space, the full extent of which is typically unknown *a priori*. Even in scenarios where conventional convergence metrics suggest adequate sampling, it remains challenging to ascertain the scale and scope of potentially unexplored regions within the broader conformational landscape where other configurational states may matter. To address this limitation, we have developed PENGUIN, a novel quantitative framework for the algorithmic comparison of observed conformational heterogeneity against a theoretical upper bound obtained from numerical simulations. The primary advantage of this approach lies in its ability to contextualize the extent of conformational sampling achieved within a given dataset size relative to a well-defined reference distribution of extreme conformational heterogeneity of equal size.

Our methodology places particular emphasis on local features at the backbone dihedral angle distribution level. We chose backbone dihedrals as our target variable for interrogating conformational distributions because ultimately the overall structural orientation of the polymer will be dictated by these distributions; however, it is important to note that there may be other collective variables worth comparing and assessing. We do not see PENGUIN as a comprehensive solution, but rather, an additional tool and framework to lend confidence in one’s simulation analysis. Importantly, we would like to make a distinction between convergence metrics at both local and global levels. Local and global conformational sampling is neither an independent nor a mutually exclusive analysis. However, while it is possible to have the convergence of global metrics, such as the radius of gyration in **Fig. S2** the convergence of global metrics is not sufficient for questions relating to local behavior - e.g., mutagenesis, post-translational modification, etc. Moreover, a lack of local conformational sampling may even, but is not guaranteed, to result in inaccurate estimations of global analyses.

While our proposed procedure for PENGUIN focuses on local backbone dihedrals, this is not the only criterion for “good” conformational sampling. Future work may extend the ideas of intra and inter trajectory analysis approach presented here to work for other collective variables or descriptors of conformational properties, such as side chain or inter-residue distances distributions. Nevertheless, we expect that comparison both within FH and between FH and EV simulations will be an informative pathway forward to help delineate folding from poorly sampled energetic traps in simulations of IDRs.

While the primary analysis and methodology here focuses on providing tooling for informing on the quality of sampling in this manuscript, the approach can easily be repurposed for the quantitative comparison of disordered protein ensembles between different reference states - e.g., wildtype and mutant, unphosphorylated and hyperphosphorylated, *etc*. Indeed, other recent approaches have used similar ideas and alternative distance metrics, such as the Wasserstein distance, have been proposed for evaluating and assessing probability distributions of conformational ensembles in disordered proteins^51^. This approach has proven useful in comparing ensembles on local and global levels; however, this metric’s time and space complexity scales quadratically with sequence length, which presents challenges for scaling this approach to many sets of simulations of large biological IDRs. Moreover, our focus here is on a special application of ensemble comparison. That is, we compare our simulations of interest to simulations of the same molecule in the excluded volume polymer limit.

By precomputing dihedral angle distributions from tripeptide simulations in the excluded volume limit for reference statistics we ensure that our tool is widely accessible for all computational biophysicists, particularly those interested in performing disordered protein simulations.

Lastly, while PENGUIN is used here as a metric to assess conformational heterogeneity compared to reference distributions, future research could explore the application of PENGUIN or its derivatives in assessing the efficacy of clustering algorithms for disordered protein ensembles. This approach holds promise, particularly in the context of Markov State Models (MSMs) constructed from extensive simulations of disordered proteins, by potentially enhancing the representativeness and accuracy of the clustering process^83,84^.

## ACKNOWLEDGEMENT

We thank members of the Holehouse lab for figure feedback and helpful discussion, as well as Kresten Lindorff-Larsen for the suggestion of investigating 2D distributions. We also thank members of the Pappu lab for many helpful discussions on sampling challenges over the years. The application of a polymer reference model is a direct extension of work by Lyle, Harmon & Pappu

## Author Contributions

JML and ASH developed the approach. JML implemented code, generated figures, acquired funding, and wrote the paper. ASH supervised research, acquired fuding, and wrote the paper.

## ABBREVIATIONS

EV: Exclude Volume
FH: Full Hamiltonian
IDR: Intrinsically Disordered Region.

## DATA AND SOFTWARE AVAILABLITY

The PENGUIN algorithm is implement in SOURSOP and can be applied directly to sets of trajectories read in there. SOURSOP is available on GitHub (https://github.com/holehouse-lab/soursop) and is installable via PypI (https://pypi.org/project/soursop/). Documentation is provided at https://soursop.readthedocs.io/ with the PENGUIN algorithm documented under the SSSampling module (https://soursop.readthedocs.io/en/latest/modules/sssampling.html). In addition, data and code used in this manuscript are provided at https://github.com/holehouse-lab/supportingdata/tree/master/2024/penguin_2024. Upon final submission, trajectories and code will also be deposited in a Zenodo repository.

