## supplementary information for "Disentangling folding from energetic traps in simulations of disordered proteins"

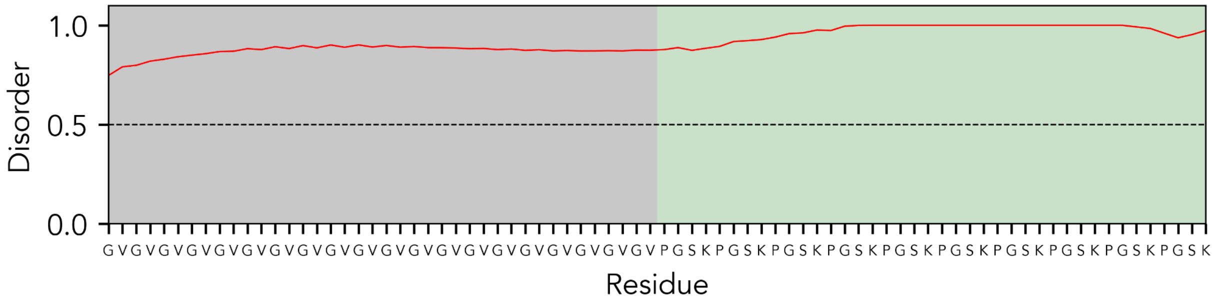


**Fig. S1:** Disorder prediction plot for the synthetic $(GV)_{20}$-$(PGSK)_{10}$ system. The gray region corresponds to the hydrophobic disordered sequence $(GV)_{20}$, which we expect to undergo nonspecific hydrophobic collapse. The green area corresponds to the sequence region we expect to sample conformational space more freely.

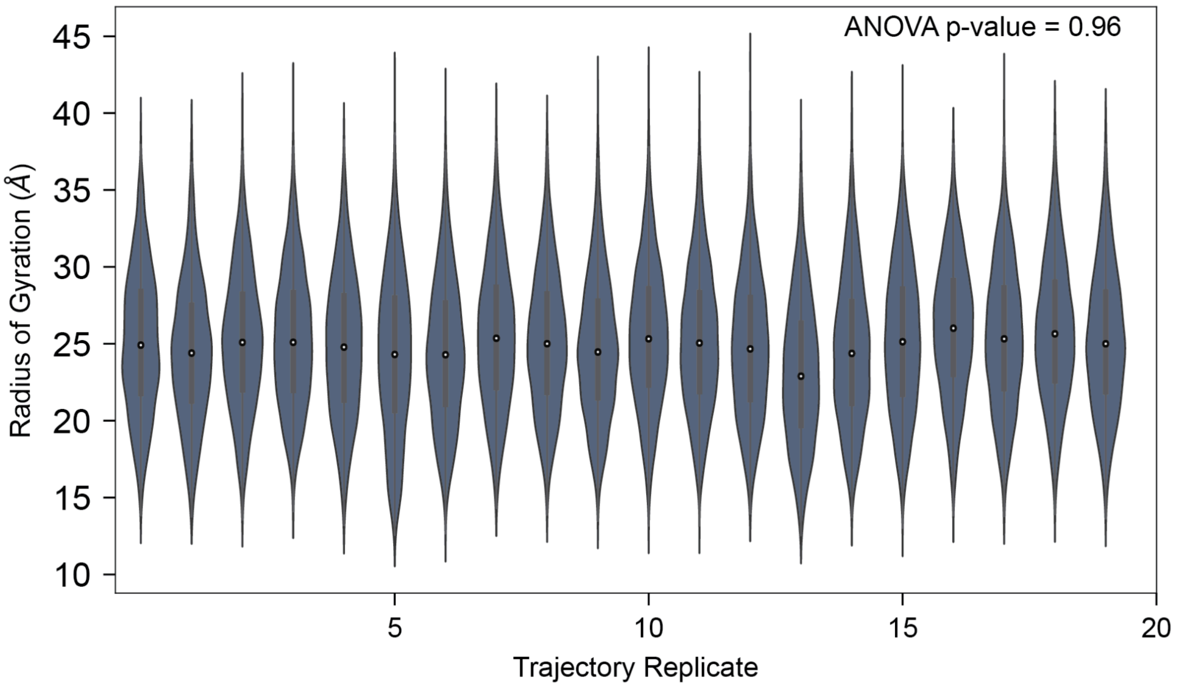


**Figure S2:** Radii of gyration distributions for the $(GV)_{20}$-$(PGSK)_{10}$ system. To determine whether the mean radii of gyration are equivalent across replicas, we performed an ANOVA test with a significance threshold of 0.05. Our analysis suggests no significant difference between the group means, suggesting the radii of gyration have converged across trajectory replicates.

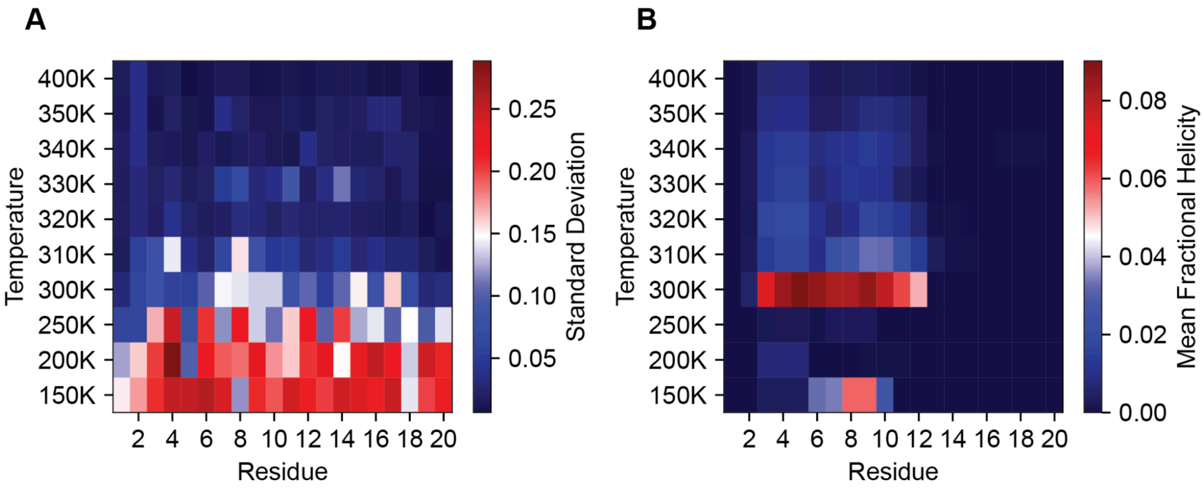


**Figure S3:** Sampling quality as a function of temperature for a transiently helical fragment of yeast GCN4. **A)** A heat map depicting the per residue standard deviation of the Hellinger distances as a function of temperature. **B)** Per residue average fractional helicity as a function of temperature. We note that while increasing temperature promotes conformational heterogeneity - as assessed by distributional similarity to the excluded volume limit - this isn't always the desired outcome, as high-temperature simulations will cause the melting of transiently structured regions.

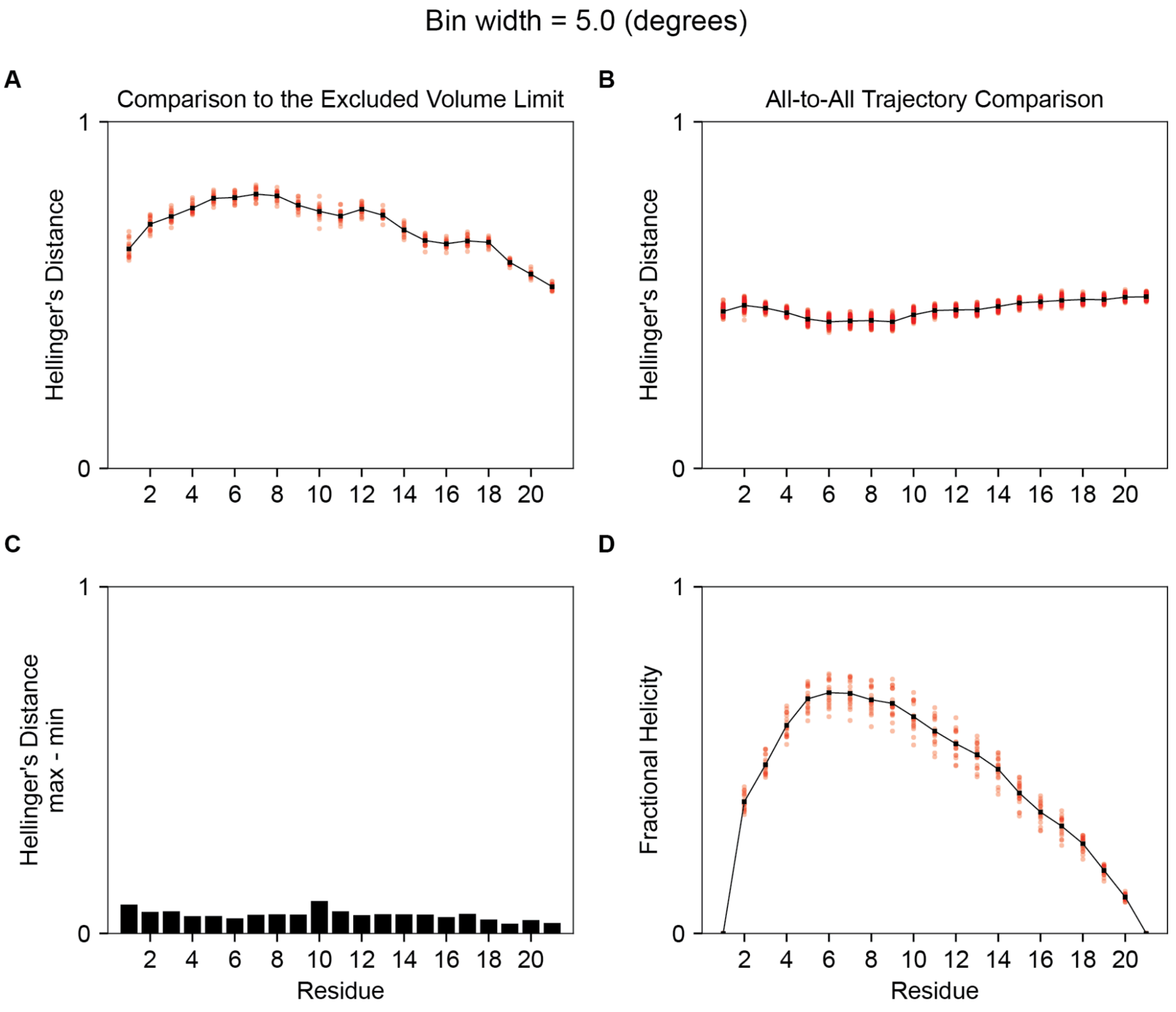


**Figure S4:** PENGUIN visualization for FS-peptide system at a bin width of 5.0 degrees. **A)** The simulated sequence **B)** Per-residue Hellinger distances between the phi dihedral angle distribution of the simulated ensemble and the excluded volume ensemble. **C)** An all-to-all simulated ensemble comparison of the per-residue phi dihedral Hellinger distance. **D)** The maximum minus the minimum Hellinger distance for each residue from the distribution in B. **E)** The per-residue fractional helicity. The transparent red points in each panel represent the raw data, whereas the black line represents the mean between replicates.

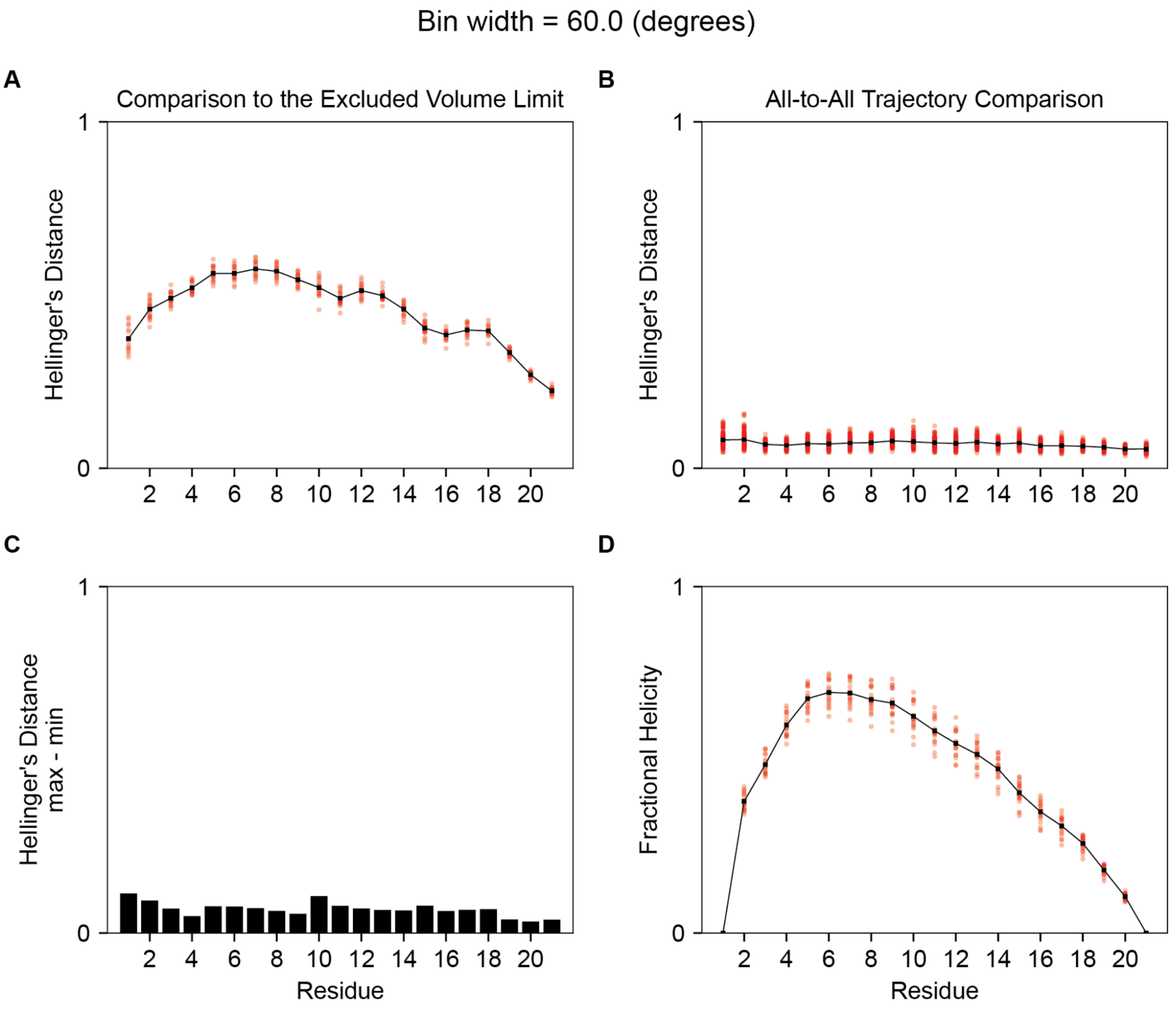


**Figure S5:** PENGUIN visualization for FS-peptide system at a bin width of 60.0 degrees. **A)** The simulated sequence **B)** Per-residue Hellinger distances between the phi dihedral angle distribution of the simulated ensemble and the excluded volume ensemble. **C)** An all-to-all simulated ensemble comparison of the per-residue phi dihedral Hellinger distance. **D)** The maximum minus the minimum Hellinger distance for each residue from the distribution in B. **E)** The per-residue fractional helicity. The transparent red points in each panel represent the raw data, whereas the black line represents the mean between replicates.

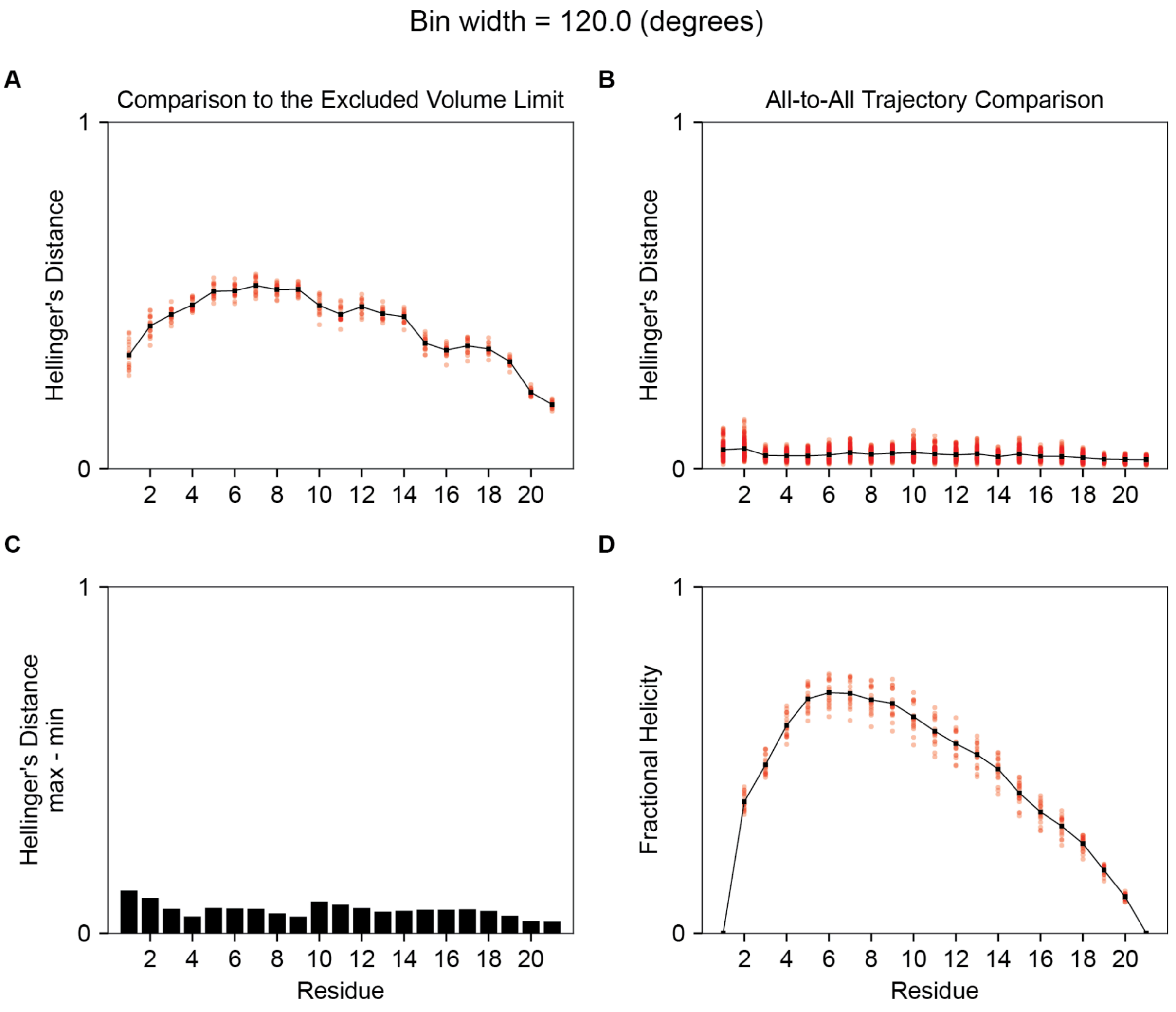


**Figure S5:** PENGUIN visualization for FS-peptide system at a bin width of 120 degrees. **A)** The simulated sequence **B)** Per-residue Hellinger distances between the phi dihedral angle distribution of the simulated ensemble and the excluded volume ensemble. **C)** An all-to-all simulated ensemble comparison of the per-residue phi dihedral Hellinger distance. **D)** The maximum minus the minimum Hellinger distance for each residue from the distribution in B. **E)** The per-residue fractional helicity. The transparent red points in each panel represent the raw data, whereas the black line represents the mean between replicates.
